# Study of combination CAR T-cell treatment for glioblastoma using mathematical modeling

**DOI:** 10.1101/2025.06.04.657886

**Authors:** Runpeng Li, Michael Barish, Margarita Gutova, Christine E. Brown, Russell C. Rockne, Heyrim Cho

**Affiliations:** Department of Mathematics, University of California Riverside, 900 University Ave., Riverside, 92521, CA, USA; Department of Stem Cell Biology and Regenerative Medicine, Beckman Research Institute, City of Hope National Medical Center, 1500 E Duarte Rd., Duarte, 91010, CA, USA; Department of Hematology & Hematopoietic Cell Transplantation and Immuno-Oncology, Beckman Research Institute, City of Hope National Medical Center, 1500 E Duarte Rd., Duarte, 91010, CA, USA; Department of Computational and Quantitative Medicine, Beckman Research Institute, City of Hope National Medical Center, 1500 E Duarte Rd., Duarte, 91010, CA, USA; School of Mathematical and Statistical Sciences, Arizona State University, 1151 S Forest Ave., Tempe, 85281, AZ, USA

**Keywords:** Immunotherapy, Chimeric antigen receptor T-cell, Mathematical oncology, Multi-scale modeling, IL13-R*α*2, HER2, EGFR, Combination therapy, CAR T antigens

## Abstract

Glioblastoma is a highly aggressive and difficult-to-treat brain cancer that resists conventional therapies. Recent advances in chimeric antigen receptor (CAR) T-cell therapy have shown promising potential for treating glioblastoma; however, achieving optimal efficacy remains challenging due to tumor antigen heterogeneity, the tumor microenvironment, and T-cell exhaustion. In this study, we developed a mathematical model of CAR T-cell therapy for glioblastoma to explore combinations of CAR T-cell treatments that take into account the spatial heterogeneity of antigen expression. Our hybrid model, created using the multicellular modeling platform PhysiCell, couples partial differential equations that describe the tumor microenvironment with agent-based models for glioblastoma and CAR T-cells. The model captures cell-to-cell interactions between the glioblastoma cells and CAR T-cells throughout treatment, focusing on three target antigens: IL-13R***α***2, HER2, and EGFR. We analyze tumor antigen expression heterogeneity informed by expression patterns identified from human tissues and investigate patient-specific combination CAR T-cell treatment strategies. Our model demonstrates that an early intervention is the most effective approach, especially in glioblastoma tumors characterized by mixed antigen expression. However, in tissues with clustered antigen patterns, we find that sequential administration with specific CAR T-cell types can achieve efficacy comparable to simultaneous administration. In addition, spatially targeted delivery of CAR T-cells to specific tumor regions with matching antigen is an effective strategy as well. Our model provides a valuable platform for developing patient-specific CAR T-cell treatment plans with the potential to optimize scheduling and locations of CAR T-cell injections based on individual antigen expression profiles.

## 1 Introduction

Chimeric Antigen Receptor (CAR) T-cell therapy is regarded as one of the most effective forms of adoptive cell-based immunotherapy for cancer treatment, with FDA approval occurring in 2017 [1, 2]. While CAR T-cell therapy has proven highly effective in treating leukemias and lymphomas [3], it is particularly well-suited for hematological cancers due to the ability of CAR T-cells to circulate through the bloodstream and lymphatic system. However, when applied to solid tumors, CAR T-cells face several challenges including: the difficulty of delivering CAR T-cells to the tumor mass, a hostile tumor microenvironment that can inhibit T-cell activity, and tumor antigen heterogeneity [3–5]. Additionally, the effectiveness of CAR T-cell therapy may be further compromised by various tumor-infiltrating cells, such as stromal cells, which support tumor growth [4].

While glioblastoma is one of the most lethal brain tumors, for decades, the standard of care has consisted of maximal surgical resection followed by a combination of radiation and chemotherapy, with the more recent addition of an electric field stimulation [6, 7]. Despite these efforts, glioblastoma remains highly aggressive and difficult to treat, with a median overall survival time of 12–18 months [8, 9] and a 5-year survival rate of 6.7%, which is the lowest among brain tumors [10, 11]. New treatment strategies for glioblastoma are actively being explored to improve clinical outcomes. Some recent advances include immunotherapy, focused ultrasound, and targeted treatments to overcome immunosuppressive tumor microenvironment [12]. Among these approaches, immunotherapy has emerged as a promising direction, aiming to reverse the immunosuppressive microenvironment and enhance the effectiveness of immune cells in recognizing and attacking glioblastoma cells. Such immunotherapies include immune checkpoint blockade, CAR T-cell therapies, oncolytic viruses and vaccines, gene therapy, and bispecific antibody therapy [13].

Mathematical models of immunotherapy for cancer have provided significant insights by predicting patient responses to treatment, identifying optimal treatment regimens, designing personalized treatment plans, and understanding the complex interactions within the tumor [14–17]. One of the first mathematical models to explain the interaction between immune cells and cancer cells is the dynamical system model developed by Kuznetsov et al. (1994) [18]. This model not only captures the population dynamics during the treatment, but also describes the formation of tumor dormant states and immune system evasion. Subsequent work by Kirschner and Panetta (1998) [19] introduced the cytokine interleukin-2 into the dynamics between tumor cells and immune effector cells. This model was able to interpret short-term tumor oscillations and long-term tumor relapses. Following this, later studies [20, 21] added periodic treatment and time delay to the model, along with corresponding stability analysis, to explain persistent oscillations observed in immune systems. Models built afterward [22, 23] incorporated additional cell types to capture tumor immune escape and explain multiple equilibrium phases of coexisting immune cells and cancer cells.

Specific mathematical models for CAR T-cell therapy include [24], a dynamical system model that investigates the correlation between the CAR T-cell dosage and the rates of proliferation and exhaustion. Kimmel et al. [25] explored the role of stochasticity in the therapy effect, offering insight into optimizing the tumor-killing rate and CAR T-cell adaptability for improved therapy outcomes. In addition, several studies have developed pharmacokinetic-pharmacodynamic (PK/PD) models for CAR-T-cells [26], demonstrating that PK/PD models not only aid in understanding the treatment, but can also guide CAR T-cell design [27]. A dynamical system model in [28] incorporates the dynamics of multiple CAR T-cells conjugates binding to cancer cells to better explain experimental results, and another model in [29] considers antigen-positive and antigen-negative tumor cells, as well as bystander CAR T-cells, to study tumor heterogeneity and bystander effects in treating solid tumors. Another recent dynamical system model in [30] integrates dual-target CAR T-cells and demonstrates its effectiveness, while providing insights into dosing regimens for dual-target CAR T-cell therapies.

To investigate the impact of spatial structure, several works have modeled the spatial-temporal interaction of CAR T-cell treatment and solid tumors, showing that spatial models can significantly contribute to designing effective treatment strategies. Fischel et al. [31] utilize a 3D agent-based model to incorporate CAR T-cell therapy, showing that the percentage of antigen-presenting tumor cells is critical in treatment success and antigen non-presenting cells can form a shield over antigen-presenting cells. A study in [32] develops a hybrid partial differential equation and agent-based model to study CAR T-cell therapy across various cancer cell lines. Luque et al. [33] develops an agent-based model to evaluate different strategies of CAR T-cell therapy for tumor-derived 3D organoids, where their findings suggest that a single dose of CAR T-cells may reduce tumor size but does not lead to complete elimination, and higher dosages can increase the number of free CAR T-cells and potential side effects. Camacho-Gomez et al. [34] presents a 3D agent-based model of CAR T-cells to compare the migratory dynamics of conventional T-cells and CAR T-cells. The authors reveal distinct migration patterns of CAR T-cells and show that a CXCL12 chemical gradient enhances the motility of CAR T-cells to be similar to conventional T-cells. In [35], the authors study resistance mechanisms of CAR T-cell immunotherapy using a system of integral-partial differential equations. By modeling continuous levels of antigen expression of cancer cells, their model enhances the understanding of relapses in CAR T-cell treatment.

Although mathematical models have been increasingly applied to investigate CAR T cell therapy, no studies have yet explored the effect of using a combination multiple CAR T-cells in glioblastoma, incorporating patient-derived tissue and spatial mathematical modeling. In this paper, we develop a 3D multiscale model to study the combination of IL-13R*α*2, HER2, and EGFR targeting CAR T-cell treatment for glioblastoma. The model combines partial differential equation (PDE) and agent-based model (ABM), using PhysiCell [36], an open-source multicellular model developing framework. In the following section, we provide the clinical motivation for using a combination of IL-13R*α*2, HER2, and EGFR targeting CAR T-cell derived from experimental data provided in [5]. Then, we investigate three treatment strategies: 1) sequential treatment, 2) multi-location spatially targeted treatment, 3) Dose-frequency-dependent treatment, and finally, propose a treatment plan that integrates our findings. Our work demonstrates that glioblastoma tissue with distinct patterns respond differently to each treatment strategies we tested, indicating the importance of personalized assessment and treatment decisions.

## 2 Results

### Selection of CAR T-cell target antigens for Glioblastoma

The study by Barish et al. [5] investigates spatial heterogeneity of antigens relevant to CAR T-cell immunotherapy in glioblastoma. This paper analyzed tumor samples from 43 patients at single-cell resolution, mapping the expression of three antigens: IL-13R*α*2, HER2, and EGFR. The findings indicate that antigen expression is not randomly distributed, but is instead clustered into distinct regions within the tumor. For example, the expression of IL-13R*α*2 and HER2 tend to be inversely related to that of EGFR, and specific patterns emerge around hypoxic areas near necrosis. This complex spatial organization may facilitate antigen escape, a phenomenon wherein the tumor cells evade CAR T-cell killing either by lack of presentation of the target antigen or by reducing antigen expression. Antigen escape is one mechanism by which treatment with single-antigen targeting CAR T-cells become less effective. The study suggests that combinatorial antigen targeting could enhance therapeutic outcomes and prevent antigen escape.

In this context, our work focuses on developing a mathematical model of glioblastoma and CAR T-cell treatment that can study different combinations of CAR T-cell treatments *in silico*. We use three tissue samples from Barish et al. [5] (PBT025, PBT018, PBT030) that display a variety of antigen expression patterns. In Figure 1, the expressions of IL-13R*α*2, HER2, and EGFR from the patient tissue samples are colored in red, green, and blue, respectively. Darker colors represent cells that express two or three antigens simultaneously, while gray represents cells that do not express any of the three antigens, considered as healthy tissue in our simulation. The three selected patient samples exhibit distinct antigen expression profiles, ranging from mixed to clustered. In particular, PBT030 primarily contains IL-13R*α*2 and EGFR antigens, which form distinct clusters in the tissue. In contrast, PBT018 contains a complex spatial mixture of IL-13R*α*2 and HER2 antigens, while also containing large regions of single antigen-positive cells. PBT025 predominantly consists of double and triple positive antigen expressing cells that are spatially mixed. Together, these three samples from [5] form a range of tumor micro-environments that could be encountered in patients.

**Fig. 1.**
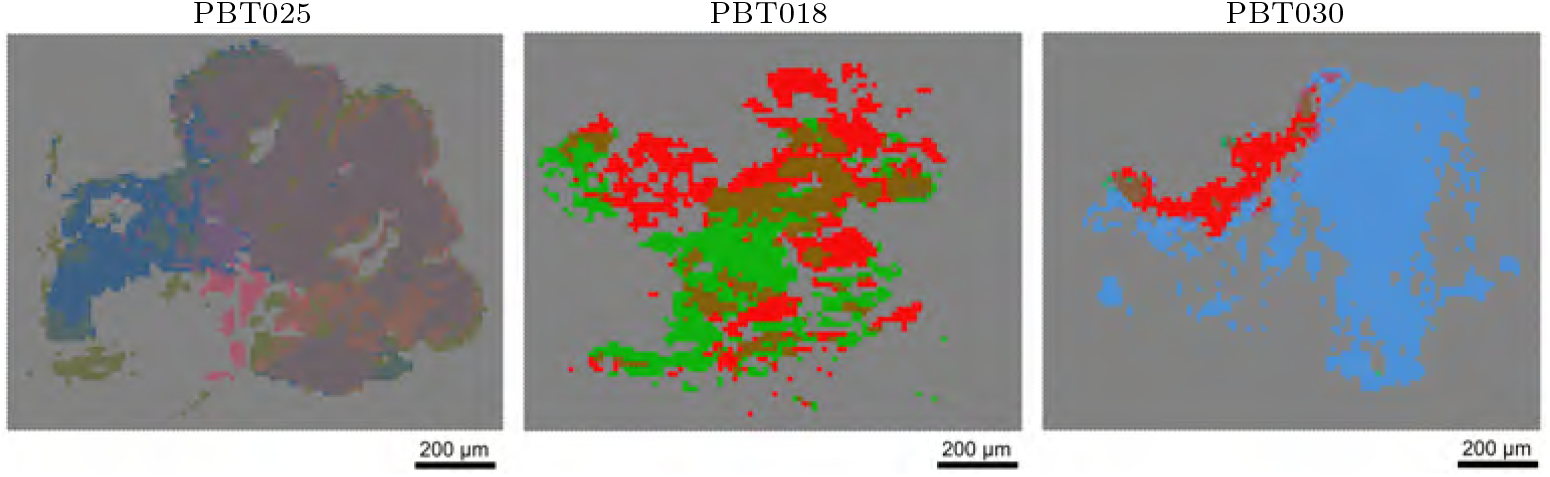
Selected glioblastoma patient tissue samples (PBT025, PBT018, PBT030) from Barish et al[5]. These tissues form the computational domain of the mathematical model. Cells are color-coded according to the type of antigen that they express, with IL-13R*α*2 (red), HER2 (green), and EGFR (blue). Mixed-colored cells represent double or triple antigen-expressing cells, and the gray cells represent cells that do not express any of the three antigens and are not considered to be targetable tumor cells in our simulation.

### Sequential treatment plan

We aim to test a sequential injection strategy motivated by the contact inhibition of locomotion observed in CAR T-cells within cancer tissue, especially when dealing with heterogeneous glioblastoma tissue with clustered patterns. For example, consider patient sample PBT018 (See Fig. 1). Suppose a biopsy at a particular location indicates presence of IL-13R*α*2-expressing glioblastoma cells. In that case, we hypothesize that administering IL-13R*α*2 CAR T-cells first, followed by HER2 CAR T-cells once the IL-13R*α*2-expressing glioblastoma cells have been cleared, could enhance the mobility and effectiveness of HER2 CAR T-cells in targeting the remaining cancer tissue. However, this strategy may be less effective in tissues with mixed or dispersed antigen-expressing patterns (e.g., PBT025) compared to those with more clustered patterns (e.g., PBT018). To assess the efficacy of sequential treatment plans, we simulate and compare simultaneous and sequential injections on the PBT018 and PBT025 tissue samples.

For the PBT018 tissue sample, we implemented two treatment plans: a simultaneous injection and a sequential injection. The simultaneous injection plan, referred to as default treatment, administers IL-13R*α*2 and HER2 CAR T-cells at the same time on day 0, while the sequential injection plan first administers IL-13R*α*2 CAR T-cells on day 0, and then injects HER2 CAR T-cells after a designated break period. We tested four different periods of break: a 5-day break, a 7-day break, a 9-day break, and an 11-day break. In both plans, we introduce 100 IL-13R*α*2 CAR T-cells and 100 HER2 CAR T-cells.

Fig. 2 illustrates the dynamics of cancer cells during the treatment period, plotting the number of cells in different subpopulations based on antigen expression. While the total number of cancer cells decreases in both plans, the sequential injection does not show any advantages over the simultaneous injection plan; however, it demonstrates similar efficacy. The percentage of tumor reduction after three weeks is 10.41% in simultaneous administration and 9.65% in sequential administration using a 7-day break. It is important to note that in the simultaneous injection plan, the HER2-expressing cancer cells do not begin to decay until around day 7. This corresponds to the time that the IL-13R*α*2 CAR T-cells eliminate the local IL-13R*α*2 cancer tissue, and the HER2 tissue is exposed. See Fig. 3 for the spatial pattern of the tissue slice. Furthermore, the number of HER2 cancer cells decays more rapidly in the sequential plan with an 11-day break, as HER2 CAR T-cells have more space to move around and find the target. Nevertheless, the simultaneous injection plan is more effective in eliminating the double-positive cancer cells expressing both IL-13R*α*2 and HER2 at the injection location than the sequential plan.

**Fig. 2.**
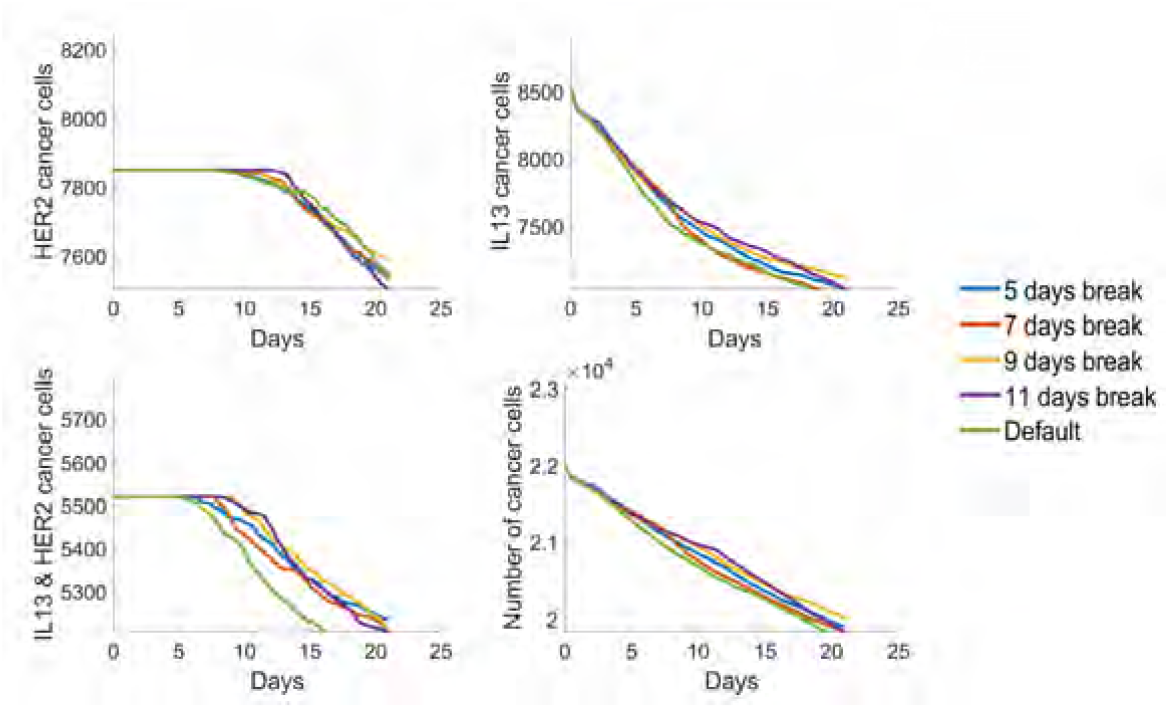
Comparison of the number of cancer cells between simultaneous (default) and sequential administration of combination CAR T-cell treatment for PBT018 tissue. In the sequential administration of IL-13R*α*2 and HER2 CAR T-cell treatments, four different time breaks are considered: 5-day, 7-day, 9-day, and 11-day breaks. The figures show the reduction of cancer cells in all categories, expressing HER2, IL-13R*α*2, and IL-13R*α*2 & HER2 antigens, including the total number of cancer cells. Considering the total number of cancer cells reduced, the sequential administration shows a similar or slightly less effective result compared to the default treatment.

**Fig. 3.**
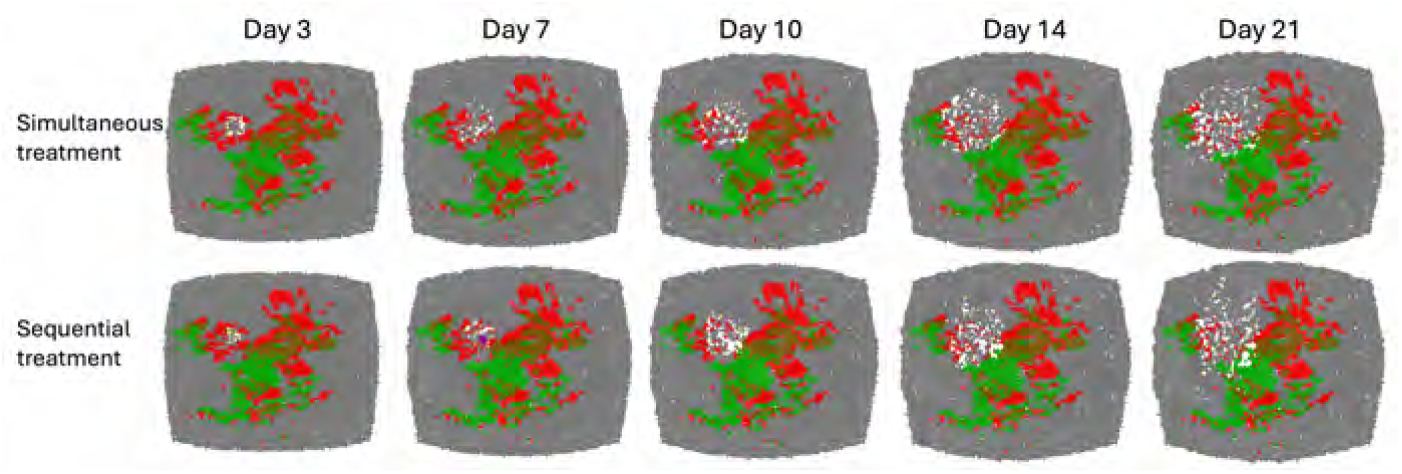
Time-course response of glioblastoma tissue sample PBT018 under combination CAR T-cell treatment, comparing simultaneous and sequential administration with a 7-day break. The tumor antigens expressed in this tissue are IL-13R*α*2 (red), HER2 (green), and IL-13R*α*2 & HER2 (brown). The simultaneous and sequential administrations show similar results considering tumor reduction (see Fig. 2) and CAR T-cell infiltration.

However, the recommendations of sequential administration may not be suitable for all patients. In particular, we propose that injecting the CAR T-cells as early as possible may be a better approach for patients with tissue that display a mixed pattern, such as PBT025. Fewer regions would restrict the CAR T-cells’ movement due to cancer cells with mismatched antigens. To test this hypothesis, we compare simultaneous and sequential administration of CAR T-cells for PBT025 patient tissue. In the simultaneous treatment, we administer 100 CAR T-cells of each type, totaling 300 CAR T-cells, injected on day 0. In the sequential treatment, we keep the total number of CAR T-cells the same; however, IL-13R*α*2 CAR T-cells are injected first on day 0, followed by HER2 CAR T-cells on day 7, and EGFR CAR T-cells on day 14. The middle slices of the simulated tissue of PBT025 as time progress are shown in Fig. 4. By comparing the snapshots taken on day 21, we observe that the simultaneous treatment has eliminated more cancer cells and spread further toward the tissue boundaries. The total number of cancer cells throughout the treatment is plotted on the right, showing that the number of cancer cells decreases more rapidly in the simultaneous treatment than in the sequential treatment starting from day 0. By the end of the treatment, the total number of cancer cells is smaller in the simultaneous treatment. Table 1 presents the percentage of tumors eliminated compared to their initial size. In PBT025, which exhibits a mixed pattern, we find that the simultaneous treatment eliminates over 14% more cancer cells than the sequential treatment by the end of the 21-day treatment. Therefore, the sequential treatment should not be considered for highly mixed tissue; instead, it is recommended to administer CAR T-cells at once as early as possible.

**Table 1.**
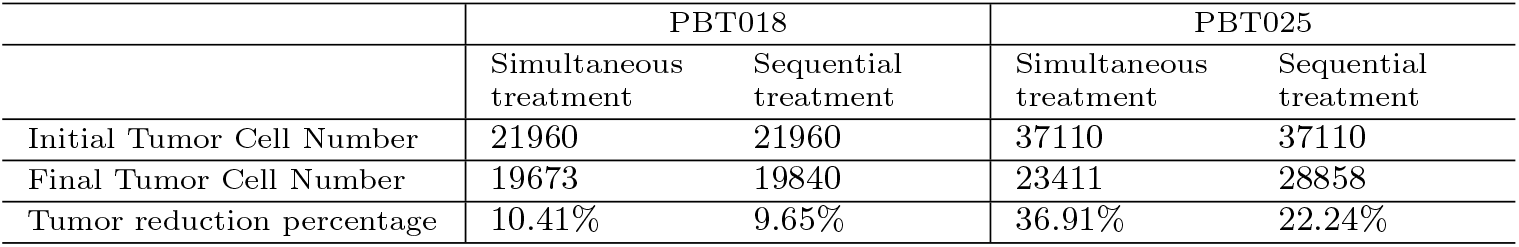
Comparison of treatment outcome between simultaneous and sequential treatment plans in PBT018 (clustered pattern) and PBT025 (mixed pattern). The simultaneous treatment is more effective in both patient tissues, but more so in PBT025 with a mixed antigen pattern. The sequential treatment gives a comparable result to simultaneous treatment only in PBT018 with an clustered antigen pattern.

**Fig. 4.**
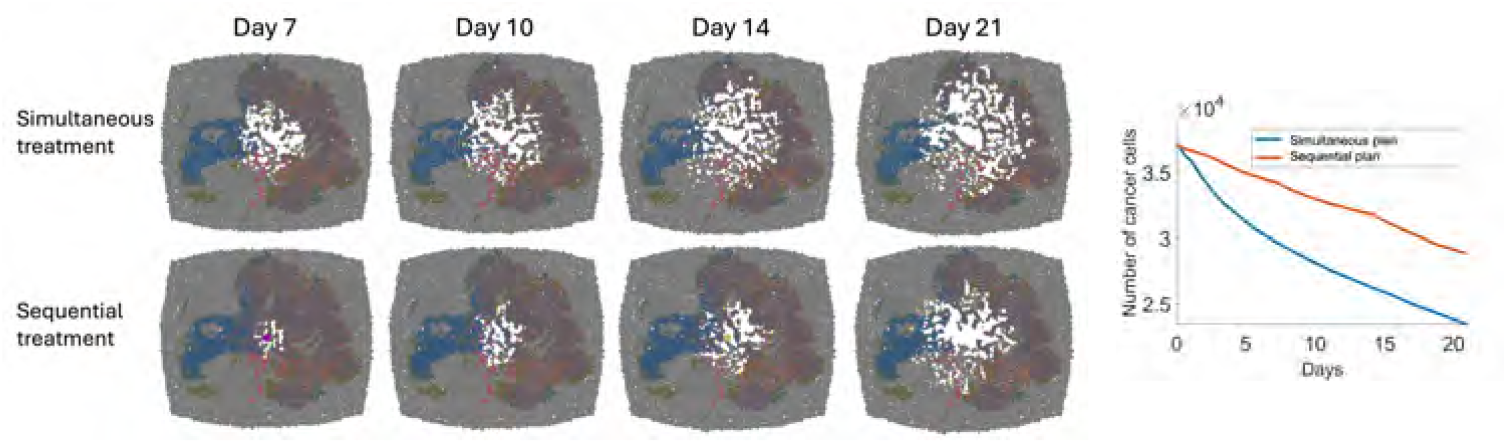
Time-course response of glioblastoma tissue sample PBT025 under combination CAR T-cell treatment, comparing simultaneous administration (Day 0) and sequential administration (Days 0, 7, and 14). The snapshots of the tissue slice (left) and the total number of cancer cells (right) are shown. The tumor antigens involved in this tissue are IL-13R*α*2 (red), EGFR (blue), and HER2 (green). The tumor cells with double and triple antigens expressed are marked in darker colors. The sequential treatment is less effective than the simultaneous treatment, considering tumor reduction and CAR T-cell infiltration.

In summary, sequential administration is not advisable compared to simultaneous administration, especially in tissue with less clustered patterns or in those with multiple areas expressing double or triple antigens. The simultaneous treatment demonstrates significantly better results for mixed tissue. However, for patients with clustered cancer tissue consisting of single antigen-expressing cells, sequential administration may be considered as a potential option, especially if spreading out the dose could offer any benefits. Nevertheless, it is important to note that simultaneous administration remains more effective in terms of cancer elimination.

### Multi-location injection plan

The second treatment strategy we aimed to investigate is a multi-location injection plan. This approach is motivated by the observation that different types of antigen-expressing cancer cells clustered in distinct locations. For instance, in sample PBT030 (See Fig. 1), IL-13R*α*2-positive and EGFR-positive glioblastoma cells are separated, with IL-13R*α*2 cells concentrated in the upper left region of the tissue. In this scenario, injecting CAR T-cells at multiple locations, specifically targeting the corresponding antigens, could lead to better treatment outcomes compared to injecting all CAR Tcells in a single location. This strategy aims to enhance the ability of CAR T-cells to identify and bind to the glioblastoma cells that express matching antigens. To test this hypothesis, we compare the efficacy of single-location and multi-location injection plans. Similar to our sequential treatment analysis in section 2, we expect treatment efficacy to differ between clustered and mixed tissue samples. Thus, we compare the two injection plans using patient tissue samples PBT030 and PBT025.

We begin by determining the injection locations, which are computed as a weighted average of the positions of cancer cells that express the same antigen. For patient tissue sample PBT030, we compute the injection locations using IL-13R*α*2 and EGFR expression levels of glioblastoma cells and use these two centers for the multi-location injection plan. In contrast, all CAR T-cells are injected at the center of the tissue for the single-location injection plan. The time progression of our simulation is shown in Fig. 5. Our results indicate that CAR T-cells in the single injection plan struggle to spread throughout the tissue, with only the EGFR CAR T-cells successfully locating and eliminating nearby cancer cells with matching antigens. On the other hand, CAR T-cells in the multi-location injection plan are more effective at removing cancer cells due to their injection at strategically chosen locations. The temporal dynamics of cancer cell reduction are shown in Fig. 5(b), demonstrating that the multi-location injection plan eliminates both IL-13R*α*2 and EGFR cancer cells more rapidly, leading to significantly better results. The overall treatment efficacy is summarized in Table 2, where the percentage of tumor reduction with the multi-location injection plan is 26.48%, nearly three times greater than the reduction achieved with the single-location injection. This outcome supports our hypothesis that treatment can be more effective when CAR T-cells are injected at locations corresponding to the relevant antigens.

**Table 2.**
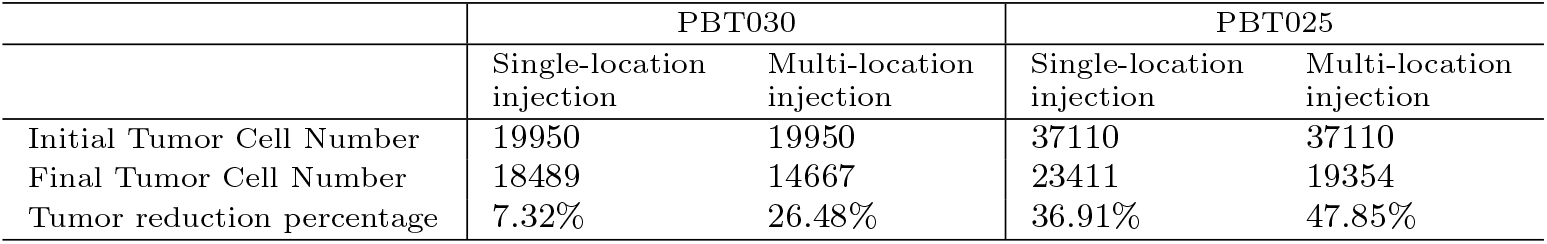
Comparison of treatment outcome between the single-location injection and multi-location injection plans in PBT030 (clustered pattern) and PBT025 (mixed pattern). The multi-location injection achieves a better result in tumor reduction than the single-location injection in both tissue samples, particularly in PBT030, which has an clustered tissue pattern.

**Fig. 5.**
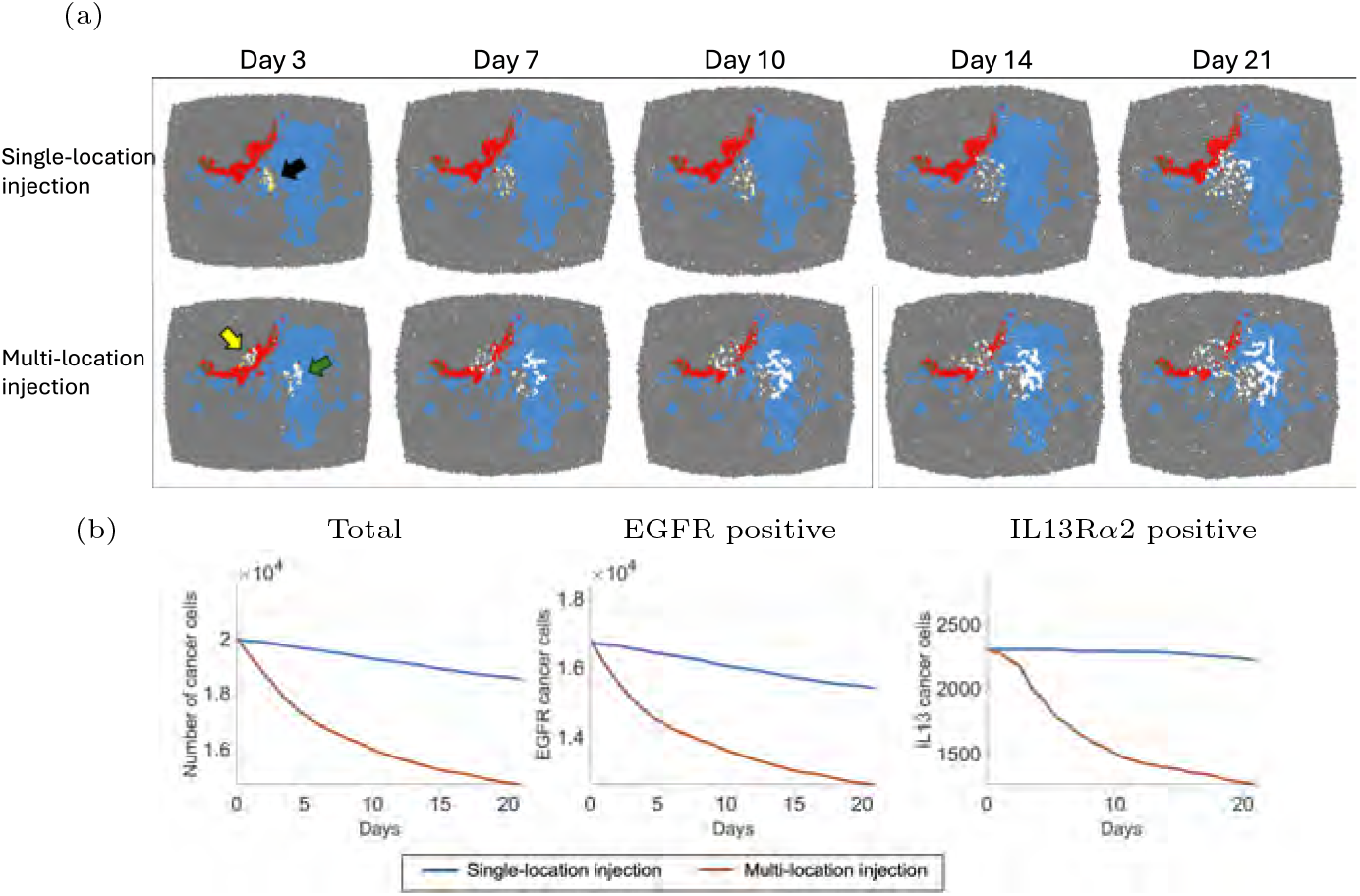
Time-course response of glioblastoma tissue sample PBT030 under combination CAR T-cell treatment, comparing single-location injection and multi-location injection plans. (a) The tumor antigens involved in this tissue are IL-13R*α*2 (red) and EGFR (blue). In the multi-location injection plan, IL-13R*α*2 CAR T-cells (yellow arrow) and EGFR CAR T-cells (green arrow) are introduced at different locations in the tumor tissue, considering the corresponding antigens. (b) The number of all cancer cells, EGFR-positive cells, and IL-13R*α*2-positive cells are shown. The multi-location injection plan reduces both EGFR and IL-13R*α*2 expressing cancer cells more rapidly than the default administration, which results in an overall improved outcome considering the tumor reduction.

With the success of the multi-location injection plan in sample PBT030, we next explore how this strategy performs in a tissue with a more mixed distribution of antigen-expressing cancer cells. We therefore simulate both treatment plans for sample PBT025 to compare their outcomes. Fig. 6 shows the time progression of tissue sample PBT025 under both single and multi-location injection plans. In this case, three antigens–HER2, EGFR, and IL-13R*α*2–are involved, and we have identified three injection locations, each corresponding to a different type of CAR T-cell. By coincidence, the injection locations for HER2 CAR T-cells and IL-13R*α*2 CAR T-cells are close. By day 21, the final snapshot of the tissue shows similar results between the two treatment plans, as the PBT025 sample contains many cancer cells expressing double and triple antigens, making it easier for any type of CAR T-cells to locate matching targets. Nevertheless, we still observe a slight reduction in cancer cell numbers under the multi-location injection plan compared to the single-location injection plan. At day 21, the percentage of tumor reduction for PBT025 tissue is 47.85% under the multi-location injection plan, compared to 36.91% with the single-location injection plan. A summary of these results, comparing PBT025 and PBT033, can be found in Table 2. While the multi-location injection shows a greater tumor reduction in the more clustered tissue sample PBT030, it also exhibits significant efficacy in the mixed tissue sample PBT025. This outcome reinforces the idea that multi-location injections are more effective than single-location injections, highlighting the importance of tailoring CAR T-cell therapies to the patient-specific heterogeneous pattern of cancer tissue.

**Fig. 6.**
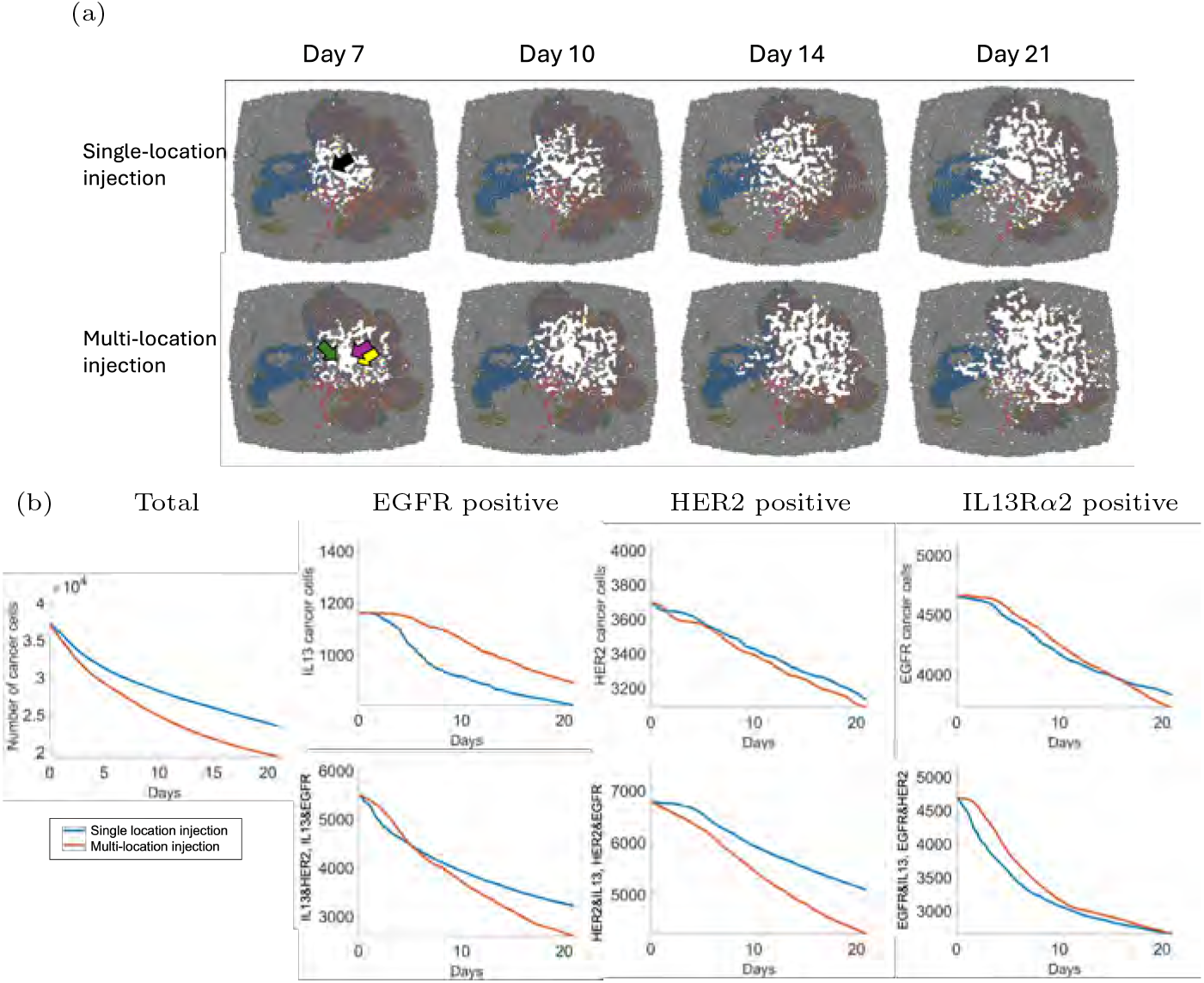
Time-course response of glioblastoma tissue sample PBT025 under combination CAR T-cell treatment, comparing single-location injection (default) and multi-location injection plans. (a) The tumor antigens involved in this tissue are IL-13R*α*2 (red), EGFR (blue), and HER2 (green). IL-13R*α*2 CAR T-cells (yellow arrow), HER2 CAR T-cells (purple arrow), and EGFR CAR T-cells (green arrow) are introduced at different locations in the tumor tissue. The results between the default and multi-location injection administrations are similar as the injection locations are similar. However, the multi-location injection shows better infiltration of CAR T-cell. (b) Considering the number of cancer cells in different cancer subpopulations, both treatment plans show similar results, while the total number of cancer cells are less in the multi-location injection plan.

### Dose-frequency-dependent treatment plan

The third treatment strategy that we aimed to investigate is a dose-frequency-dependent injection of CAR T-cells, focusing on how dosage affects treatment efficacy. We use a fixed total number of CAR T-cells and compare different injection schedules. For example, we administer larger doses less frequently, while smaller doses are given more often, maintaining the total number of CAR T-cells. We apply this strategy to the patient tissue sample PBT025, where CAR T-cells are introduced at the center of the tissue. Each antigen type receives a dosage of 180 CAR T-cells, summing up to a total of 540 CAR T-cells. We compare three injection plans: high-frequency, medium-frequency, and low-frequency. In the high-frequency plan, we inject 30 CAR T-cells of each antigen type (a total of 90 CAR T-cells) per dose, administering six doses every 3 days. For the medium-frequency and low-frequency treatment, we inject 60 and 90 CAR T-cells of each antigen type (a total of 180 and 270 CAR T-cells), with three doses every 7 days and two doses every 14 days, respectively. The simulation runs for 21 days.

Fig. 7 shows the progression of the tissue slice and the total number of cancer cells over time using the three dose-frequency-dependent treatment plans for PBT025. The low-frequency treatment demonstrates better efficacy than the other two plans, especially in the early stages when the CAR T-cells are initially injected. We observe a correlation between CAR T-cell dosage and tumor tissue infiltration by CAR T-cells. This is further evidenced by the trend in the cancer cells, where the slope of tumor reduction is steeper with higher doses. The percentage of tumor reduction, presented in Table 3, indicates that the low-frequency treatment can eliminate 5% more tumor cells than the other two plans. This finding is consistent with the established observation that earlier administration of more CAR T-cells lead to more effective tumor elimination. However, while beneficial, the difference in tumor size is modest–ranging from 30% to 35% over the 21-day treatment period.

**Table 3.** Comparison of treatment outcome between three treatment plans with different doses and frequencies in tissue sample PBT025. The low-frequency plan results in a moderately improved tumor reduction compared to the other plans.

|  | PBT025 |  |  |
| --- | --- | --- | --- |
|  | High-frequency | Medium-frequency | Low-frequency |
| Initial Tumor Cell Number | 37110 | 37110 | 37110 |
| Final Tumor Cell Number | 25941 | 25630 | 24065 |
| Tumor reduction percentage | 30.1% | 30.94% | 35.15% |

**Fig. 7.**
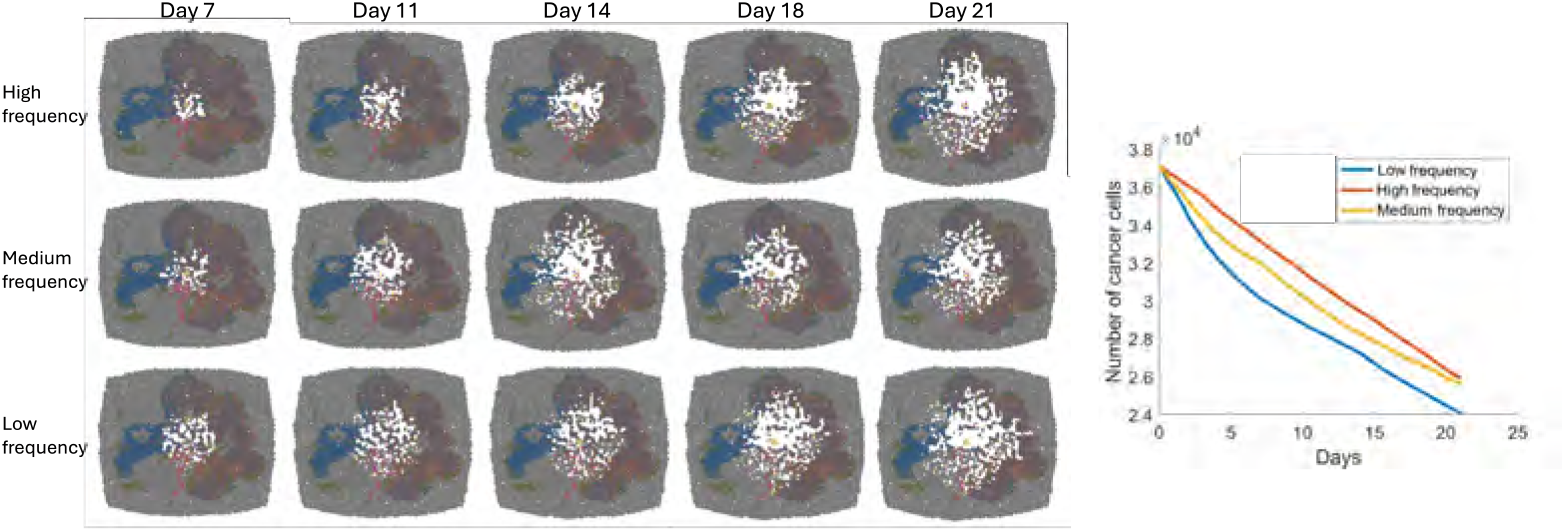
Time-course response of glioblastoma tissue sample PBT025 under combination CAR T-cell treatment, studying dose-frequency-dependent administration. The snapshots of the tissue slice (left) and the total number of cancer cells (right) are shown. The IL-13R*α*2, EGFR, and HER2 CAR T-cells are introduced at the center with three different doses (90, 180, and 270 CAR T-cells from top to bottom) with corresponding time intervals (3, 7, and 14 days, respectively). The treatment plan with the strongest dose shows the most tumor reduction, especially at the beginning when the CAR T-cells are first injected.

### Combination treatment

So far, we have explored three different treatment strategies based on timing, injection location, and dose frequency. In this section, we will combine the key insights from the previous simulations: 1) multi-location injections can be effective in both clustered and mixed cancer tissues, and 2) larger doses administered at lower frequencies may yield better outcomes.

Based on these observations, we aimed to integrate these two strategies and evaluate their long-term efficacy. We will compare two treatment plans: a weekly injection plan with CAR T-cells injected at the center (Treatment Plan 1) and a bi-weekly double-dose plan with multi-location injection (Treatment Plan 2). Both plans are simulated on tissue samples PBT025 and PBT030. CAR T-cells are injected over the first 8 weeks, followed by an additional 8 weeks of monitoring after the final dose. The treatment Plan 1 involves eight doses of 25 CAR T-cells for each type, while treatment Plan 2 consists of four doses of 50 CAR T-cells for each antigen type. Both injection plans administer the same total dose of CAR T-cells, which is, 600 CAR T-cells. The multi-location injection of treatment Plan 2 are determined as the weighted center of each subpopulation, as in section 2, while treatment Plan 1 has all CAR T-cells injected at the center.

In Fig. 8, we observed that treatment Plan 2 eliminates more cancer cells than treatment Plan 1 for PBT025, resulting in less remaining cancer tissue visible in the tissue slices. The trajectory of total cancer cell numbers shows that treatment Plan 2 consistently outperforms treatment Plan 1 throughout the treatment. After 16 weeks, treatment Plan 2 eliminates approximately 14% more cancer cells than treatment Plan 1. A similar result can also be seen in PBT030, where treatment Plan 2 eliminates 16% more cancer cells than treatment Plan 1. A summary of these results is provided in Table 4. This simulation confirms that combining multi-location injection with a higher-dose-lower-frequency plan is more effective than the default treatment Plan 1 for both PBT025 and PBT030 patient tissues.

**Table 4.** Comparison between the default weekly treatment (Plan 1) and multi-location injection coupled with bi-weekly higher dose treatment (Plan 2) in PBT025 and PBT030. The treatment Plan 2 enhances tumor reduction by approximately 15% compared to Plan 1 by the end of the treatment period.

|  | PBT025 |  | PBT030 |  |
| --- | --- | --- | --- | --- |
|  | Plan 1 | Plan 2 | Plan 1 | Plan 2 |
| Initial Tumor Cell Number | 37110 | 37110 | 19950 | 19950 |
| Final Tumor Cell Number | 12048 | 7023 | 7381 | 4172 |
| Tumor reduction percentage | 67.53% | 81.08% | 63.00% | 79.09% |

**Fig. 8.**
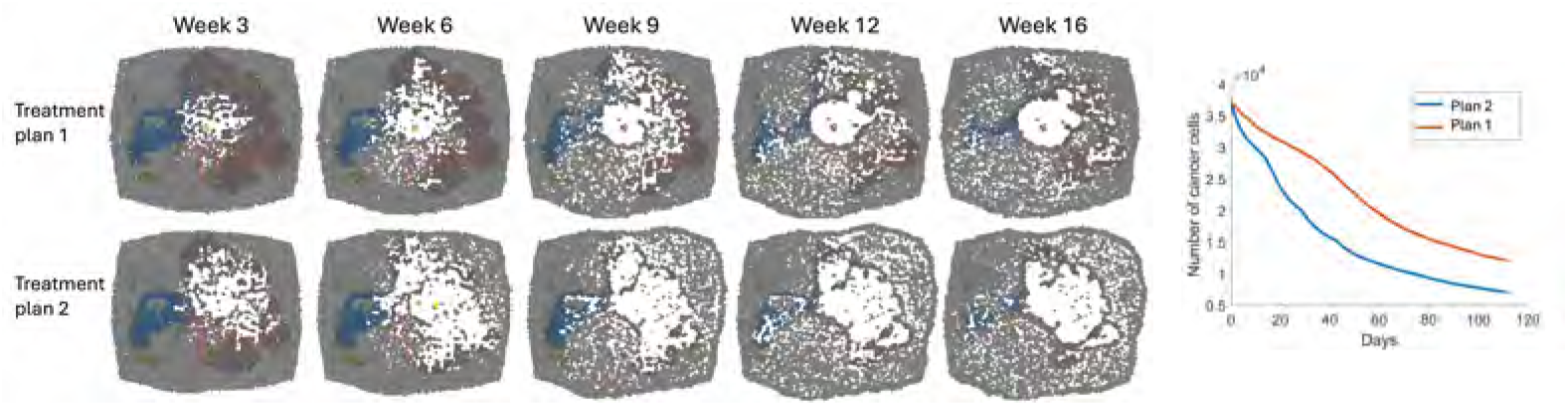
Time-course response of glioblastoma tissue sample PBT025 under combination CAR T-cell treatment, comparing treatment Plan 1 (weekly default) and Plan 2 (biweekly, multi-location injection) in the long term. The snapshots of the tissue slice (left) and the total number of cancer cells (right) are shown. The treatment Plan 2 demonstrates superior outcomes in tumor reduction and CAR T-cell infiltration compared to Plan 1.

## 3 Discussion

In this paper, we develop a mathematical model to study the combination CAR T-cell treatments for glioblastoma. Our model is built using PhysiCell, a platform to develop multiscale models of interacting cells within a dynamic 3D microenvironment [36]. By modeling the heterogeneous population of glioblastoma cells and multiple CAR T-cell populations with a cell-based model, our model incorporates the heterogeneity of glioblastoma tissue of individual patients to optimize CAR T-cell treatment. We modeled glioblastoma cells that can express any combination of three different antigens – IL-13R*α*2, HER2, and EGFR – expressed at a continuous level. The three implemented antigens cover most of the glioblastoma tissue introduced in [5]. Thus our model can be used to study combination CAR T-cell treatments for general glioblastoma patients.

Using our model, we explored three different treatment strategies – timing, injection location, and dose/frequency – for combination CAR T-cell treatments, comparing these approaches to a default treatment, which involves a single injection of all CAR T-cells at the center of the tissue. From the simulation results, we conclude three observations: i) sequential treatment can be significantly less effective in mixed tissue; however, it can be comparable in clustered tissue, ii) multi-location injection treatment tends to be effective across all tested tissues, but to a greater degree in clustered tissue, and iii) low-frequency treatment is more effective than treatments with lower doses and higher frequencies. In summary, injecting CAR T-cells targeting specific locations of matching antigens can significantly enhance treatment efficacy. Additionally, larger doses can effectively eliminate more cancer cells even with longer intervals between each injection. Notably, patients whose tumors exhibit more clustered tissue patterns of antigen expression show more pronounced variability in treatment response, indicating that treatment decisions for such patients require greater precision and individualized assessment. Moreover, for such patients, sequential injection plan may be a viable option, as it allows for the spreading out the doses, that may help reduce patients’ side effects, such as cytokine storm syndrome and immune effector cell-associated neurotoxicity syndrome [37, 38].

Despite the effort that has been made in this work, several important questions remain to be addressed. An imminent future work is to calibrate the model to additional clinical data, in particular, data that captures distinct killing efficiency of IL-13R*α*2, HER2, and EGFR CAR T-cells [39–41]. Currently, the model is calibrated to *in vitro* experiment data of IL-13R*α*2 CAR T-cell treatment only [24]. Further experimental data on HER2 and EGFR CAR T-cells will allow for more accurate model calibration and predictions. In addition, the tumor microenvironment is highly complex and involves multiple interacting components [42]. To better capture this complexity, we plan to improve the model by incorporating various subpopulations of CAR T-cells, such as regulatory T-cells and exhausted T-cells, and details of the microenvironment, including cytokines. These enhancements will enable a more comprehensive analysis of treatment dynamics and potentially guide more effective therapeutic strategies. Additionally, the study of dosage and frequency should consider patients’ side effects, such as cytokine storm syndrome and immune effector cell-associated neurotoxicity syndrome [37, 38]. Although our findings suggest that larger doses with longer injection intervals can result in smaller tumor sizes, it is crucial to investigate the maximum doses that patients can tolerate, taking potential adverse reactions into account. We acknowledge another limitation of our work that is the scale discrepancy between our simulation and actual glioblastoma tissue. Our study includes 80,000 cells, which is relatively small compared to the actual cell numbers presented in real patient tumor, and the simulated CAR T-cell dosage is correspondingly low. We proposed to increase the size of the simulation and include various scenarios, such as surgery before the administration of CAR T-cell therapy. Furthermore, since our multi-location injection study indicates the importance of determining the prioritized treatment area, we plan to explore additional practical approaches, such as combining intratumoral and intraventricular delivery [39, 43].

## 4 Methods

We develop a hybrid partial differential equation and agent-based model of CAR T-cell therapy for glioblastoma based on the open-source multicellular code, PhysiCell [36]. The model describes the dynamics and interactions of cells in a three-dimensional microenvironment, with cell phenotypes dependent on the dynamic changes of the tumor microenvironment. It incorporates essential cell behaviors, such as cell cycling, cell death, volume regulation, motility, and cell-cell mechanical interactions. The model is coupled with the bio-transport solver, BioFVM [44], which efficiently simulates reaction-diffusion PDEs of environmental substrates in 3D.

In our ABM model, we consider four distinct cell types: glioblastoma cells, and three types of CAR T-cells that target the antigens IL-13R*α*2, HER2, and EGFR. The PDE model of the tumor microenvironment includes two biochemical substrates: oxygen and an immunostimulatory factor, such as IL-2 for example. Glioblastoma cells and CAR T-cells are modeled using off-lattice agents, with cell volume computed using a dynamical system model that represents changes in liquid and solid fractions within the cell. Each cell can exist in one of three states: proliferating, quiescent, or necrotic—depending on the local oxygen concentration, and the model accommodates cell cycle progression, division, and death accordingly. Cell migration is influenced by cell-cell adhesion, cell-cell repulsion, chemotaxis, and random motility. Glioblastoma cells secrete the immunostimulatory factor, which attract CAR T-cells to migrate along its gradient. The immunostimulatory factor and oxygen are modeled using reaction-diffusion equations, and these are coupled to the cells in the ABM through the reaction term. The detailed model equations, adapted from the default PhysiCell implementation, are provided in the Appendix A.

### ABM interaction model between glioblastoma and CAR T-cells

In the ABM, the glioblastoma cells exhibit specific phenotypes characterized by the expression levels of antigens EGFR, IL-13R*α*2, and HER2. We denote the expression levels as ***a*** = (*a*_1_, *a*_2_, *a*_3_) *∈* [0, 1]^3^. A glioblastoma cell is considered to be positive for the *i*-th antigen if *a*_*i*_ is over some threshold, denoted as *a*_*thr*_.

The three types of CAR T-cells can form an adhesion and subsequently eliminate the glioblastoma cell if the matching antigen level *a*_*i*_ is over the threshold *a*_*thr*_. The adhesion between the glioblastoma cells and CAR T-cells is determined by a probability that is linearly dependent on the antigen expression levels as well as the distance between the cells. The adhesion probability is defined as follows:

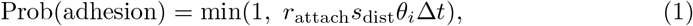

where *r*_attach_ is the CAR T-cell’s rate of forming new cell adhesions, *s*_dist_ is the scaled distance between the target glioblastoma cell and the attaching CAR T-cell, and *θ*_*i*_ is the scaled expression level of the *i*-th antigen on the glioblastoma cell defined as follows:

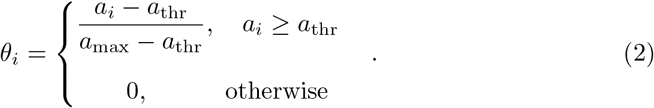

The maximum level of antigens *a*_max_ = 1 represents the maximum value of the antigen expression level from the data. The scaled distance is computed as follows:

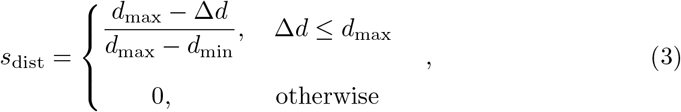

where *d*_max_ and *d*_min_ denote the maximum and minimum adhesion distance, and Δ*d* denotes the distance between the glioblastoma and CAR T-cell. According to this setup, the adhesion between the glioblastoma cell and *i* type CAR T-cell is formed only if the distance between two cells is less than *d*_max_ and the antigen level *a*_*i*_ exceeds the threshold *a*_thr_. We assume that multiple CAR T-cells can attach to a single glioblastoma cell, and each event is independent.

When two cells are attached, the CAR T-cells attempt to induce apoptosis in the attached glioblastoma cell with the following probability,

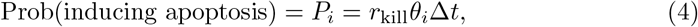

where *r*_kill_ is the killing rate. Here, we assume that apoptosis probability depends linearly on the antigen expression level. If multiple types of CAR T-cells are attached, each CAR T-cell attempts to induce apoptosis independently. Thus, the overall probability of a glioblastoma cell being eliminated can be computed by using the survival probability *P*_survive_ = Π_*i∈I*_ (1 − *P*_*i*_), where *I* is the set of attached CAR T-cell antigen index among {1, 2, 3}, and it follows that the overall probability of successful cancer cell elimination is *P*_kill_ = 1 − *P*_survive_.

The scheme of the ABM model for the glioblastoma and CAR T-cell interaction are as follows. First, we determine if a CAR T-cell is attached to any glioblastoma cell. If it is attached, the CAR T-cell will attempt to induce apoptosis with a probability given by Eq. (4) and if it is successful, the CAR T-cell proceeds with the killing process. If apoptosis is not induced, it will detach with a probability Prob(detach) = Δ*t/*(*T*_attach_ + 10^−15^), where *T*_attach_ is the duration of the attachment. On the other hand, if the CAR T-cell is not attached to a glioblastoma cell, it will search for surrounding cells to attach to with a probability given by Eq. (1). In this case, the CAR T-cell will check all glioblastoma cells within the maximum adhesion distance. A summary flowchart of the interaction model is in the Appendix A.

### Parameter calibration

We aim to calibrate the model to the experimental data presented in [24], which includes *in vitro* data of IL-13R*α*2 CAR T-cell treatment of glioblastoma cells. Given the large number of parameters in the model, we first conduct a parameter sensitivity analysis on several key factors in the model: oxygen uptake rate, oxygen decay rate, adhesion strength, repulsion strength, maximum adhesion distance, oxygen diffusion coefficient, CAR T-cell attachment rate to glioblastoma cells, and CAR T-cell killing rate of glioblastoma cells. We find that the attachment rate *r*_attach_ and the killing rate *r*_kill_ are the most critical parameters affecting tumor apoptosis. Consequently, we focus on calibrating these two parameters to the data from [24]. The optimal parameters are identified by minimizing the residual sum of squares between the model’s predictions and the experimental data of tumor volume. For the calibration, we consider a range for attachment rate *r*_attach_ *∈* [0.5, 5] and the killing rate *r*_kill_ *∈* [0.1, 0.2]. The experiments in [24] examine glioblastoma cells mixed with IL-13R*α*2-targeted CAR T-cells at different ratios-1:5, 1:10, and 1:20-and with varying antigen expression levels (low, medium, and high). In our simulations, we assign the antigen expression levels *a*_*i*_ to 0.35, 0.65, and 0.95 for low, medium, and high antigen-expressing glioblastoma cells, respectively. The experimental data and the best model fit are shown in Fig. 9, where we find the optimal parameters to be *r*_attach_ = 2 and *r*_kill_ = 0.15. These values were determined through a grid-based search with a grid size of 0.5 for the attachment rate and 0.1 for the killing rate. The values for the remaining model parameters are estimated from various biological literature sources. Due to the lack of specific data for HER2 and EGFR-targeted CAR T-cells, we assume that the attachment and killing rates for all three types of CAR T-cells (IL-13R*α*2, HER2, and EGFR) are identical for the purpose of this study.

**Fig. 9.**
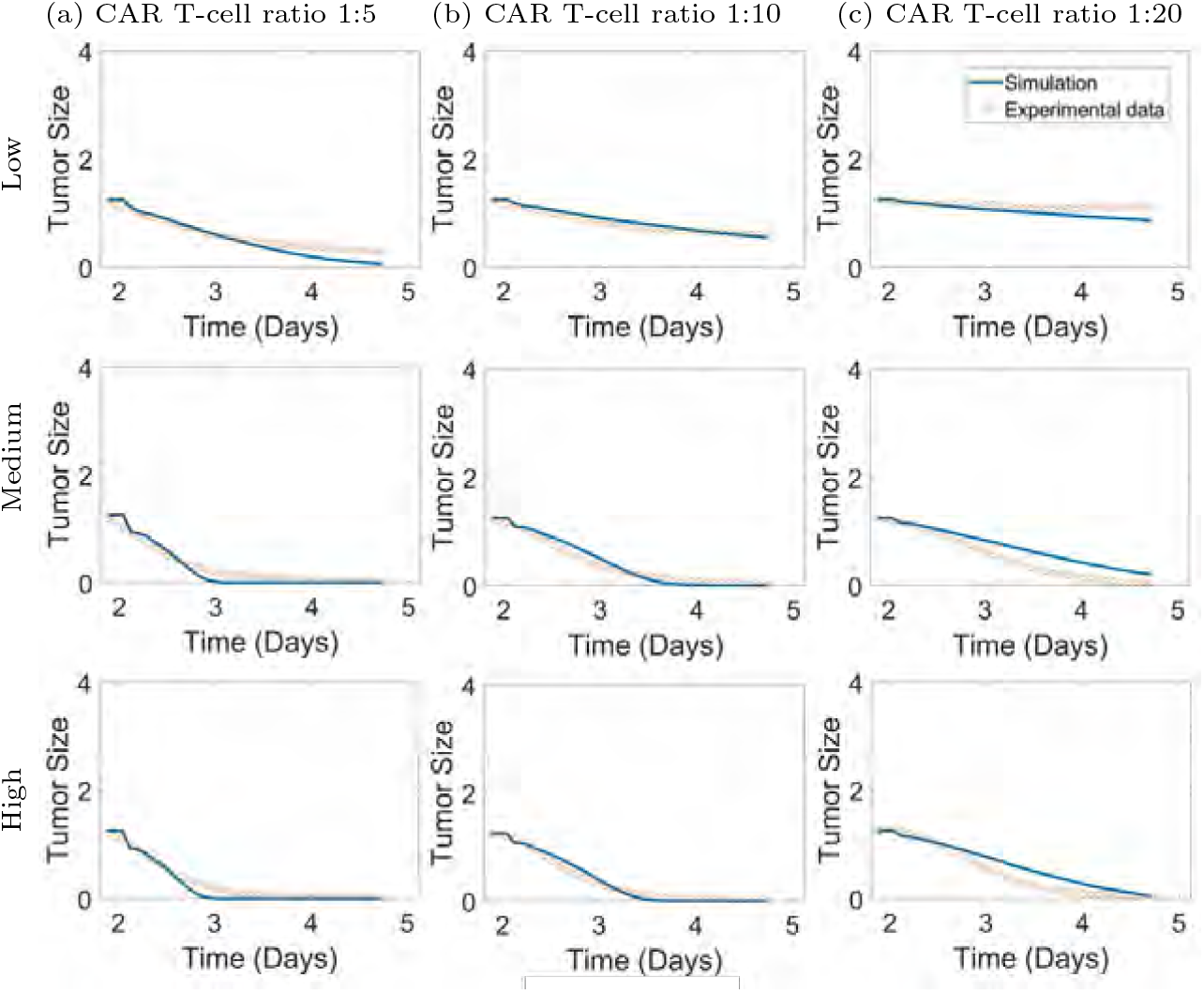
The optimal model calibration using the attachment rate *r*_attach_ = 2.0 and the killing rate *r*_kill_ = 0.15. The experimental data (circle) collected in [24] and the calibrated model simulation are shown. Columns (a), (b), and (c) represent CAR T-cell to cancer cell ratios of 1:5, 1:10, and 1:20, respectively. Rows indicate cancer cell antigen density levels from low to high.

## Acknowledgements

Research reported in this publication was supported in part by the City of Hope Biostatistics and Mathematical Oncology shared resources supported by the National Cancer Institute of the National Institutes of Health under grant numbers P30CA033572. The content is solely the responsibility of the authors and does not necessarily represent the official views of the National Institutes of Health.

## Declarations

### Data availability

All of the data supporting the results of this study are available within the paper and its Supplementary Information. The raw, de-identified patient data are available from the corresponding authors upon reasonable request and subject to Institutional Review Board approval.

### Code availability

The code will be available in https://github.com/tony1747/ upon publication.

### Author contribution

H.C., R.R., M.B., M.G. conceptualized and designed the study. R.L. and H.C. developed the model, and R.L. developed the code and performed the analyses. All authors contributed to the interpretation of results. R.L. and H.C. drafted the manuscript, and all authors revised and approved the manuscript. H.C. supervised the overall direction and planning of the study.

## Appendix A Model equations of PDE/ABM based on Physicell

The computational domain is 1800 *×* 1500 *×* 160 microns initialized with approximately 80000 glioblastoma cells. As the patient tissue sample is provided in 2D as shown in Fig. 1, we construct a 3D, prism-like volume by vertically stacking 10 identical copies of the cross-section.

### A.1 PDE system of microenvironment substrates

The biochemical environment is modeled as a system of reaction-diffusion PDEs for two chemical substrates: oxygen and an immunostimulatory factor, denoted as ***ρ*** = (*ρ*_1_, *ρ*_2_) = (*ρ*_1_(***x***, *t*), *ρ*_2_(***x***, *t*)). The equation is of the form

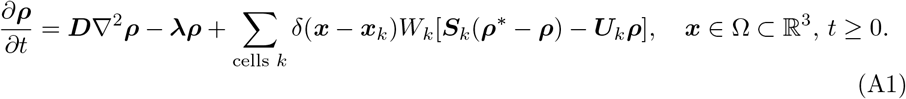

Here, *δ*(***x***) is the Dirac delta function, ***x***_*k*_ = ***x***_*k*_(*t*) is the position of the *k*-th cell, *W*_*k*_ is the *k*-th cell’s volume, ***S***_*k*_ is the *k*-th cell’s source rates, ***U***_*k*_ is the *k*-th cell’s uptake rates, ***ρ***^*\**^ is the saturation densities, ***D*** is the diffusion coefficients, and ***λ*** is the decay rates. We assume Dirichlet boundary condition for oxygen and zero flux boundary condition for the immunostimulatory factor on *∂*Ω. The cell volume *W*_*k*_ = *W*_*k*_(*t*) is updated through an ODE system of that describes the changes in liquid and solid fractions within the cell. The glioblastoma and CAR T-cells uptake oxygen and the glioblastoma cells secrets the immunostimulatory factor. The parameter values are taken to be identical to the cancer immunology example in [36].

### A.2 Cell migration model in ABM

The velocity of a cell agent is calculated based on cell-to-cell repulsion/adhesion, and cell motility as

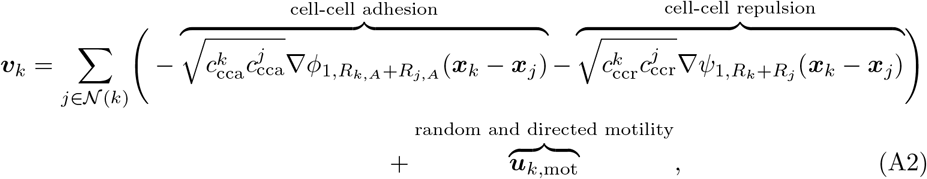

where 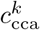 and 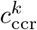 are the *k*-th cell’s cell-cell adhesion and repulsion parameters, *R*_*k*_ is its radius, *R*_*k,A*_ is its maximum adhesion distance, and *ϕ*_*n,R*_(***x***) and *ψ*_*n,R*_(***x***) are adhesion interaction and repulsion interaction potential functions, respectively, defined as

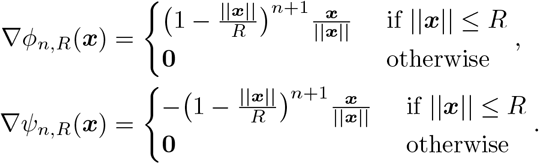

The cell motility of the *k*-th cell ***u***_*k*,mot_, is given by:

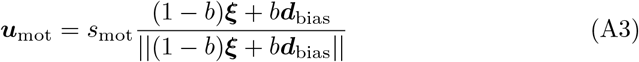

where *s*_mot_ is the migration speed, ***ξ*** is a random unit vector for random motion, and ***d***_bias_ is a unit vector for directed motion, and *b* is a weight parameter that determines the fraction between the random and directed migration. For the CAR T-cells, the directed migration is updated through

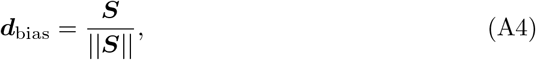

where ***S*** = *∇ρ*_2_(***x***_*k*_(*t*), *t*) is the gradient of the local immunostimulatory factor.

### A.3 Flowchart of caner and CAR T-cell interaction model in ABM

The flowchart in Figure A1 summarizes the interaction model of glioblastoma and CAR T-cells. See section 4 for the detail of probabilistic models describing each events.

**Fig. A1.**
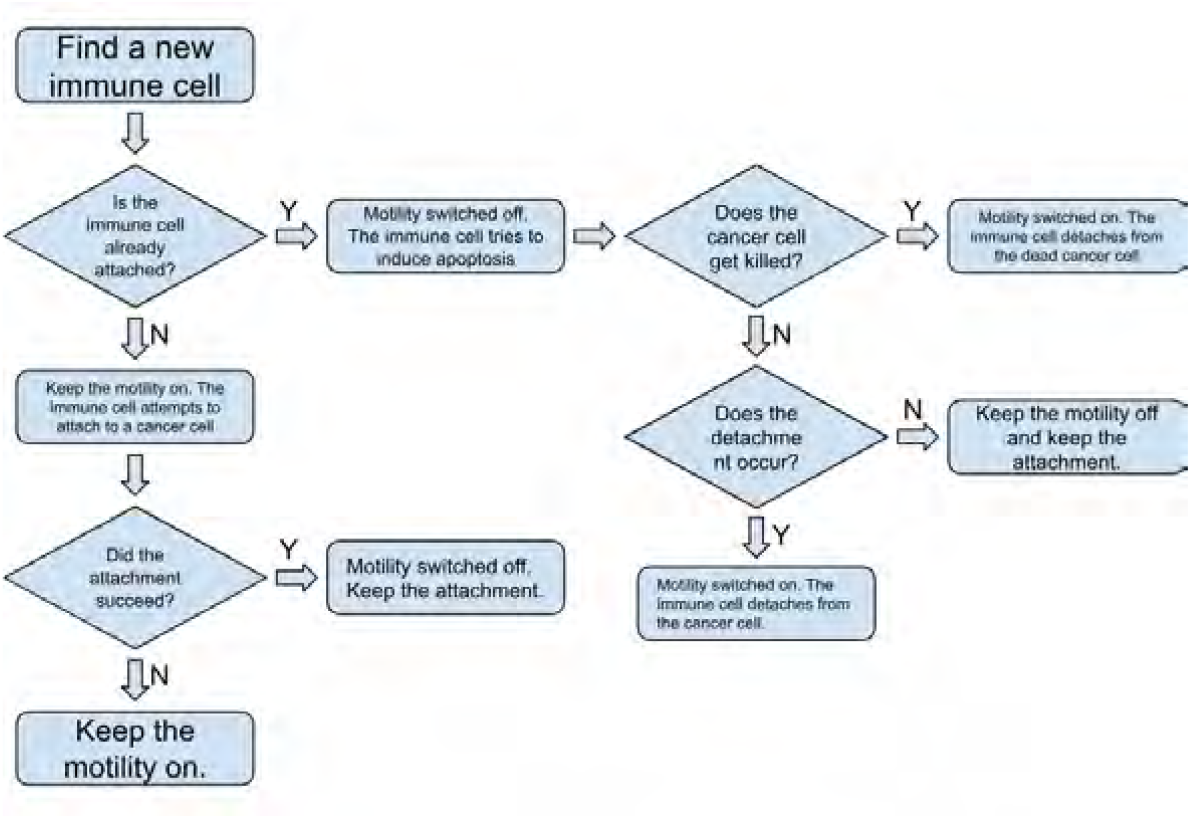
Summary of the CAR T-cell and cancer cell interaction process in the ABM.

